# Structural Characterization of Cardiac Purkinje Fibers Using Inhomogeneous Magnetization Transfer (ihMT): A proof of Concept MRI-Histology Approach

**DOI:** 10.64898/2026.01.29.702467

**Authors:** Arash Forodighasemabadi, Evgenios N. Kornaropoulos, Marion Constantin, Lucas Soustelle, Fanny Vaillant, Jude Leury, Richard Walton, Olivier Bernus, Bruno Quesson, Olivier M. Girard, Guillaume Duhamel, Julie Magat

## Abstract

**BACKGROUND:** The cardiac Purkinje network plays a critical role in maintaining synchronized activation of ventricles but remains challenging to image due to its fine and unique structure. Conventional MRI techniques lack sufficient contrast to distinguish the underlying structural composition of Purkinje Fibers (PF).

**PURPOSE:** This study investigates the potential of inhomogeneous Magnetization Transfer (ihMT) as a novel contrast mechanism for visualizing and differentiating subregions of the PF.

**METHODS:** Five fixed ex-vivo sheep hearts (n = 5) containing free running PF were scanned with a 9.4T MRI using a 2D ihMT RARE sequence.

**ASSESSMENT:** ihMTR maps were analyzed using manually defined regions-of-interest (ROIs) corresponding to free-running PF, insertion points, and myocardium. Histological analysis (light and polarized microscopy) was performed on matched sections to quantify collagen types I and III, adipocytes, Purkinje cells, and cardiomyocytes.

**RESULTS:** Three ihMT protocols, which produced high ihMTR values in free-running PF (9.25–10.83%) and strong absolute contrast relative to the myocardium (2.00–2.17%) and insertion points (2.99–3.40%) in one sample were selected and applied to all samples. Across all samples, mean ihMTR in free-running was consistently higher than in insertion points (11.5 ± 1.5% vs. 9.0 ± 2.9%). Histological analysis revealed a significantly greater collagen content in free-running regions compared with insertion points (72.4 ± 15.9% vs. 31.1 ± 13.1%; p = 0.001), along with higher adipocyte content at insertion points vs. free-running regions (12.3 ± 6.1% vs. 3.8 ± 2.7%, non-significant). Collagen type III was more prominent at insertion points but remained a minor component overall.

**CONCLUSION:** ihMT imaging can distinguish PF subregions based on microstructural differences, particularly collagen and adipocyte distribution. This study lays the groundwork for developing biophysical models to interpret ihMT signals in terms of tissue composition and microstructure, providing a foundation for future studies.

**Sponsor:** This study received financial support from the French Government by the National Research Agency (ANR; SYNATRA ANR-21-CE19-0014-01) and Région Nouvelle Aquitaine (convention N°AAPR2022-2021-16609210).

## Introduction

The His-Purkinje system is a specialized network of fibers that plays a critical role in the synchronous activation of the ventricles by providing a rapid conduction pathway ^1^. This coordination ensures efficient ventricular contraction, which is essential for maintaining proper cardiac function ^2^. However, the Purkinje system is also implicated in the initiation and maintenance of ventricular fibrillation, which can lead to sudden cardiac death ^3^. Understanding this intricate system is therefore crucial for preventing and treating cardiac arrhythmias ^4^.

From a structural point of view, Purkinje cells, the cellular component of the His-Purkinje network, are considerably larger than myocardial cells in their transverse section size ^5^. They also exhibit morphological differences across species; for instance, ungulates have large, clearly differentiated Purkinje Fibers (PF) with few peripheral myofibrils and abundant central glycogen, while rodents possess PFs that are structurally very similar to working myocardial cells ^6^. In ungulates like sheep and pigs, PFs within false tendons are surrounded by a dense collagen sheath. This sheath is thought to act as an electrical insulator and likely contributes to the rapid and efficient conduction of the action potential and may also provide mechanical support and protection to these specialized conducting cells ^2,6^. There are also structural differences within the PF itself, notably between free-running part surrounded by collagen and optimized for rapid conduction and insertion points, which are responsible for transmitting impulses to the slower-conducting myocardial tissue ^7^.

Recent advancements in imaging technologies have significantly improved the study of the His-Purkinje system. Micro-computed tomography (micro-CT), enhanced with iodine-based contrast agents, has facilitated high-resolution visualization of the cardiac conduction system (CCS) ^8–10^. For instance, Stephenson et al. used contrast-enhanced micro-CT to distinguish major subdivisions of the CCS in rat and rabbit hearts, including the Purkinje network ^11^. Later, this method was employed to generate the first 3D representation of the human cardiac conduction system ^8^ and to examine morphological changes in pathological hearts ^10^. Similarly, Aminu et al. utilized micro-CT with graphene oxide contrast agents to reconstruct the CCS in fixed human hearts ^9^ and Chen et al. used contrast-enhanced CT to study anatomical changes in Myocardial Infarction ^12^.

In contrast to the CT, Magnetic resonance imaging (MRI) is a non-ionizing modality which can provide 2D and 3D images in any orientation, with contrasts that can be adjusted depending on the acquisition technique. However, the required spatial resolution to visualize the 3D morphology of conduction pathways remains a major challenge. Hwang et al. used high-resolution MRI to trace conduction pathways in isolated rabbit hearts, demonstrating its capability to map the Purkinje-ventricular junctions in 3D ^13^. However, conventional MRI techniques, based on relaxation properties (T_1_, T_2_, and T_2_^*^) and proton density, often lack sufficient contrast for effectively differentiating PF from the surrounding cardiac muscle ^14^. An ex-vivo study on swine hearts has shown that rotating frame relaxation maps can provide contrast between the atrioventricular conduction axis and myocardium, demonstrating the potential of quantitative MRI techniques for characterizing the conduction system ^15^.

Magnetization Transfer (MT) has demonstrated sensitivity to highly organized macromolecules, including collagen. Several studies on phantoms ^16^ and biological tissues such as intervertebral discs ^17^, knee cartilage ^16^, renal ^18^ and liver fibrosis ^19^, and myocardial scars ^20^ have shown that the MT signal correlates with collagen content. Magat et al. ^14^ explored MT contrast for imaging the Purkinje network and provided optimized sequence parameters to acquire 3D images of the purkinje fibers in ex-vivo samples of porcine heart.

Inhomogeneous Magnetization Transfer (ihMT) ^21–24^ is an extension of MT and is sensitive to dipolar order relaxation ^23^. IhMT has been primarily applied to brain ^22,25^ and spinal cord ^26^ imaging due to its high sensitivity and specificity to myelinated tissue ^27^, making it particularly useful for studying demyelinating pathologies such as Multiple Sclerosis ^28^. Given the sensitivity of ihMT for tissues with highly organized structure and the known correlation between MT signal and collagen content, applying ihMT to cardiac conduction system particularly PF, which are rich in collagen, may hold potential for improved characterization of these structures. In this study, we present an experimental exploration of ihMT sequence parameters aimed at enhancing contrast for improved visualization of Purkinje fibers in ex vivo cardiac samples. MR imaging findings are complemented by histological analysis to support tissue characterization.

## Materials & Methods

### Experimental setup

Five samples (S1-5), all taken from the left ventricle and containing myocardium along with free-running Purkinje fibers (Figure 1-a), were obtained from the hearts of five female sheep (age= 9±0.7 y, weight = 50.5 ± 6.5 kg) following euthanasia via sternal thoracotomy under general anesthesia. The animal protocol has been described in Cabanis et al. ^29^. The tissues were fixed in 4% formaldehyde containing 0.1% gadoterate meglumine (0.5 mmol/mL; Dotarem, Guerbet, France). The animal protocol was approved by the Animal Research Ethics Committee (CEEA50) in accordance with the European rules for animal experimentation (European legislation 2010/63 EU). For MRI data acquisition, each sample (approximate size of S1: 35 × 35 × 25 mm^3^, S2: 55 × 35 × 35 mm^3^, S3: 30 × 25 × 35 mm^3^, S4: 35 × 30 × 30 mm^3^, S5: 35 × 25 × 30 mm^3^) was placed in a syringe filled with Fluorinert (a liquid that does not produce a signal in proton MR; CF2, Sigma-Aldrich, St. Louis, MO) (Figure 1-a) to mitigate potential susceptibility artifacts. The syringe was equipped with a temperature probe (SA Instruments, NY) connected to a computer for continuous monitoring of temperature during the experiments (Figure 1-a). The samples were heated using water bath tubing wrapped around the syringe.

**Figure 1:**
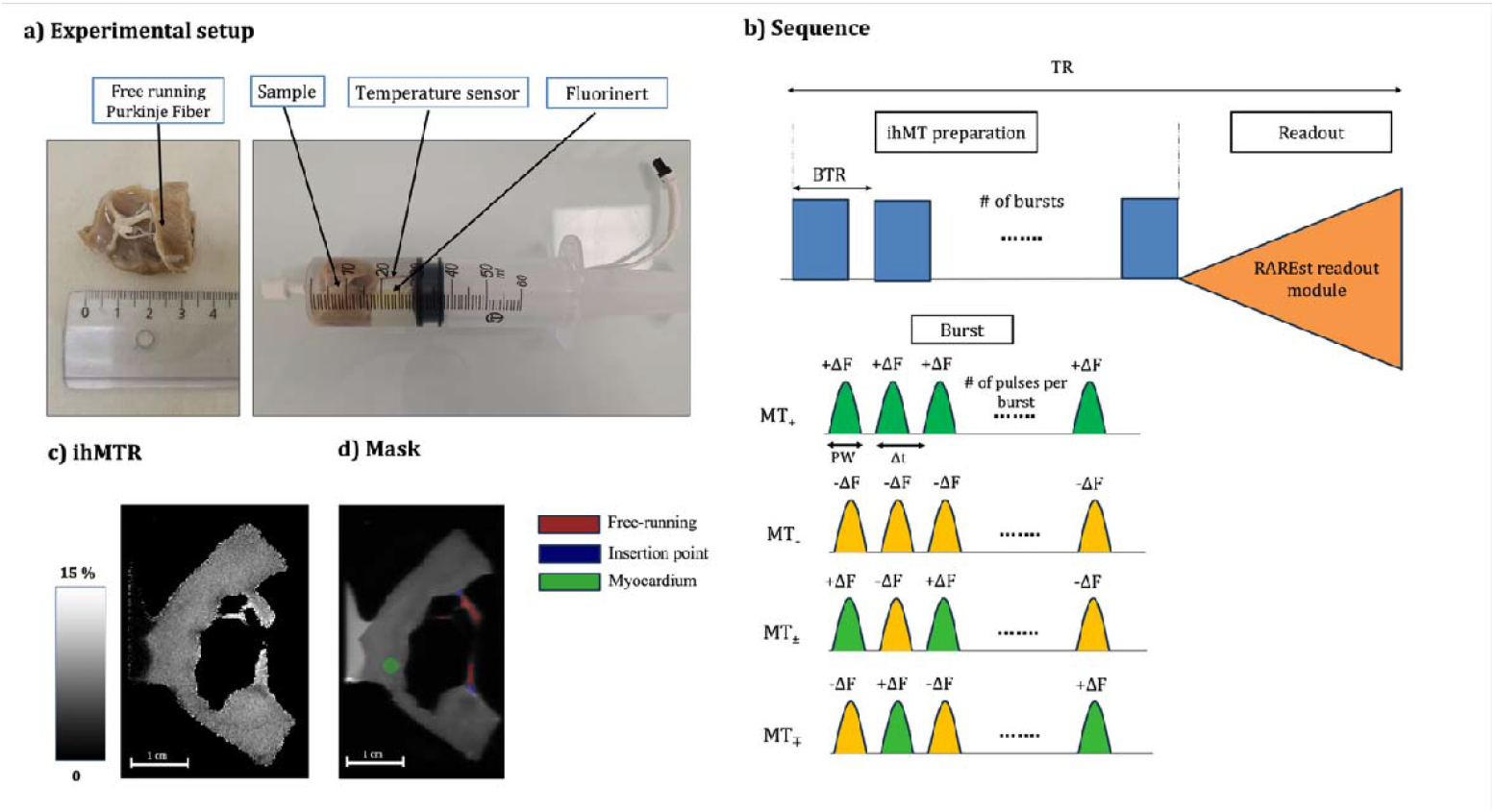
a) Experimental setup showing the sample containing myocardium and PF placed in a syringe filled with Fluorinert, equipped with a temperature probe connected to an external computer for monitoring. b) 2D ihMTRARE sequence, featuring ihMT preparation with bursts of MT pulses followed by readout (PW: Pulse Width; Δt: repetition delay; BTR: Burst TR). c) an example of ihMTR map in % d) The mask on M_0_ including free-running PF (red), Insertion point (blue) and myocardium (green).

In the first phase of the study, termed *Protocol selection*, one sample was heated to 33□±□1□°C and subjected to a range of ihMT preparation saturation parameters to identify optimal contrast settings. During the ihMT preparation, parameters such as pulse width, B_1_ amplitude, number of pulses, and offset frequency (Figure 1-b) were systematically varied, as they are known to significantly influence ihMT signal and contrast. In the second phase, termed *Protocol application*, three optimized protocols selected from the first phase were applied on all five samples, heated to 36 ± 1°C.

### MR acquisition

All MR experiments were performed at 9.4 T using a BioSpec 94/30 scanner (Bruker BioSpin MRI, Ettlingen, Germany) equipped with a cylindrical transmit coil (200-mm internal diameter) and a 4-channel phased-array receive coil and equipped with gradients with a maximum 300 mT/m strength.

After acquiring the localizers, GRE sequences (in axial, coronal and sagittal views) were run to identify the best slice for visualizing both the PF and the myocardium. A 2D ihMT-prepared centric-out single-shot RARE sequence (Figure 1-b) was used. The ihMT preparation consisted in the acquisition of 4 MT-weighted images obtained different saturation preparations. These include positive single-sided (MT_+_), positive-negative dual-sided (MT_±_), negative single-sided (MT_-_) and negative-positive dual-sided (MT_∓_) images. Dual-sided saturations were implemented using frequency-alternated radiofrequency (RF) pulses that alternate between positive (+Δf) and negative (-Δf) frequency offsets.

Since the diameter of the free-running PF can vary and can go up to more than a millimeter, Partial Fourier acquisition (factor□=□1.8) was used, enabling the collection of 200 volumes of each of the 4 MT weighted image within a total scan time of approximately 2 hours. For thin fibers in the free-running region, Partial Fourier was omitted due to increased partial volume effects and blurring, and the number of MT-weighted volumes was reduced to 50 per image to maintain the same acquisition time. Other parameters included a TE of 20 ms, a TR of 3000 ms, 10 averages of reference images without any preparation module (MT_0_), and a voxel size of 0.25 × 0.25 × 1mm^3^.

### MR images post processing

IhMT ratios (ihMTR, expressed in %) were calculated as:

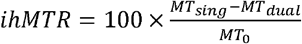

with MT_sing_=MT_+_+MT_-_ and MT_dual_=MT_±_+MT_∓_. IhMTR maps were generated using the publicly available script: https://github.com/lsoustelle/ihmt_proc(hash f3f49e0)). An example of ihMTR map is shown in Figure 1-c.

Three regions of interest (ROIs) were manually drawn using the ITK-SNAP toolbox (www.itksnap.org) ^30^. One ROI was placed over the Free-running segment of the fiber (Figure 1-d, red), another at the Insertion point where the fiber enters the myocardium (Figure 1-d, blue), and a third within the Myocardium (Figure 1-d, green).

In the *Protocol selection* phase of the study, MT saturation parameters detailed in Table 1 were investigated on sample 1. The contrast values (in ihMTR units) were defined as:

**Table 1:**
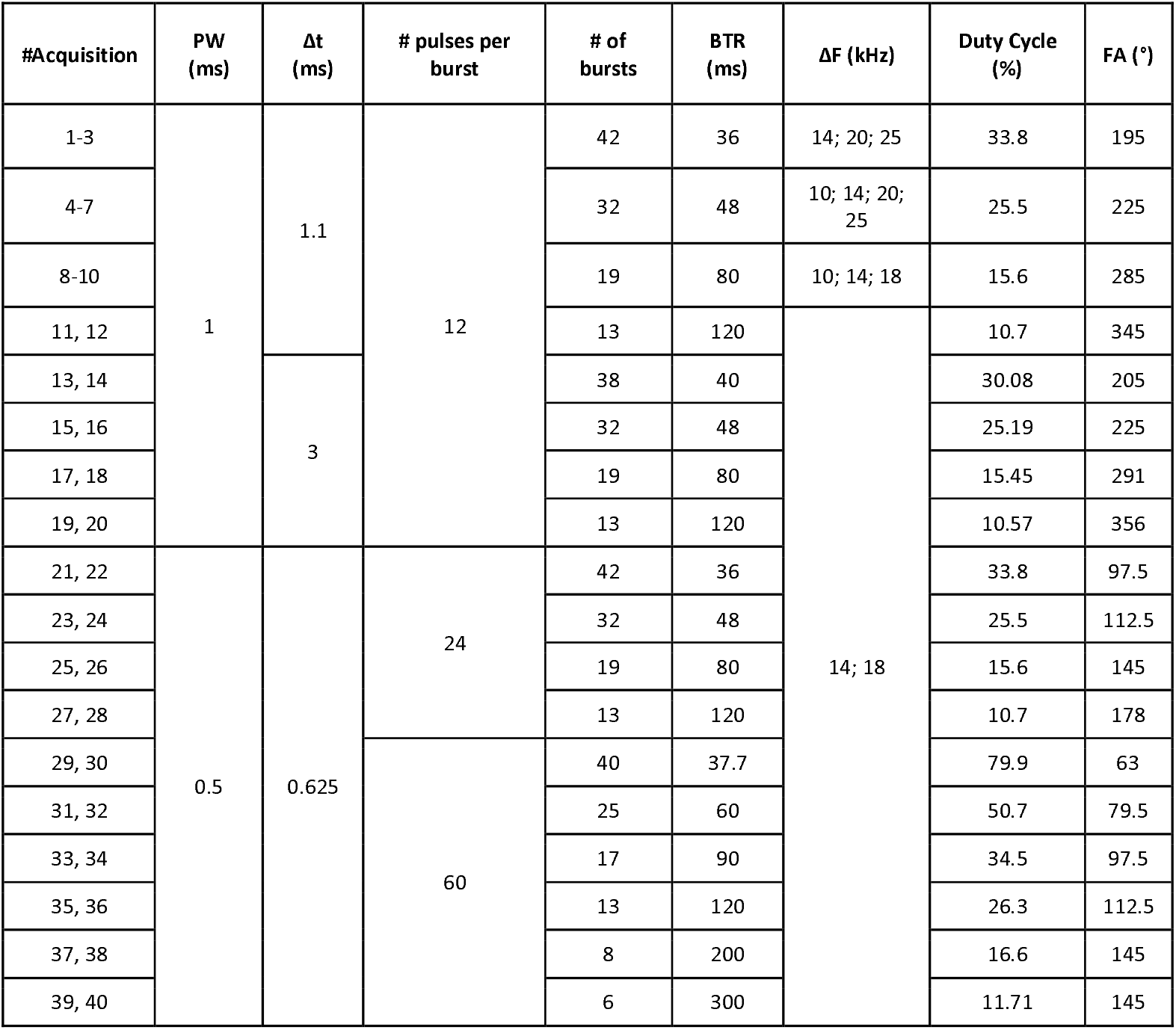
All ihMT sequence configurations were applied with Hann-shaped pulses for a saturation B1RMS (defined over BTR) maintained constant at 9 ± 0.3 μT. As an example, based on the table, parameters for acquisition #12 is: PW/Δt/#pulses per burst/ #bursts/ΔF=1 ms/1.1 ms/12/13/120 ms/18 kHz and acquisition #13 is: PW/Δt/#pulses per burst/ #bursts/ΔF=1 ms/3 ms/12/38/40 ms/14 kHz (PW: Pulse Width; Δt: repetition delay; BTR: Burst TR).

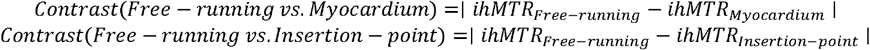

Based on the results obtained, three protocols listed in Table 2 were selected and were used in the *Protocol application* phase on all five samples.

**Table 2:** The three selected protocols providing the best contrast on sample 1 during the *Protocol selection* phase (PW: Pulse Width; Δt: repetition delay; BTR: Burst TR).

| Protocol | PW (ms) | $\Delta t$ (ms) | # pulses per burst | # Bursts | BTR (ms) | $\Delta F$ (kHz) |
| --- | --- | --- | --- | --- | --- | --- |
| Pr1 | 1 | 1.1 | 12 | 32 | 48 | 20 |
| Pr2 | 0.5 | 0.625 | 24 | 32 | 48 | 18 |
| Pr3 | 0.5 | 0.625 | 60 | 17 | 90 | 18 |

### Histological analysis

After MR experiments, histological analyses were conducted to provide detailed insights into the tissue structure. After dehydration, samples were embedded in paraffin and sectioned at 6 μm. For samples S1–S5, 2, 2, 3, 4, and 2 microscopic slides, respectively, were selected as the best anatomical matches to the corresponding ihMTR imaging slices. Tissue sections were stained with Picro Sirius Red for structural identification: collagen fibers, myocytes, and adipocytes were red, yellow and white, respectively. The polarized light (PL) filter (analyzer-polarizer, Nikon) allowed for differentiation between collagen type I (in red, orange and yellow) and collagen type III (in green) ^31^. Slices were examined at 20X magnification on a tissue slide scanner (Axio Scan Z1, Carl Zeiss Microscopy GmbH, Jena, Germany) or at 4X magnification optical microscope (Eclipse 80i, Nikon, Tokyo, Japan).

#### Histological images post processing

Segmentation of adipocytes and collagen in Picro Sirius Red histology images was performed using ImageJ with manual color thresholding. The segmentation process involved adjusting hue, saturation, and lightness to isolate blue pixels corresponding to collagen and dark pixels corresponding to adipocytes. This approach enabled the creation of binary masks for both tissue components.

For collagen segmentation under polarized light, a home-made Python program was written to convert RGB images (Red-Green-Blue in PNG format) color space to HLS (Hue-Lightness-Saturation) to facilitate color-based classification ^32^. The hue values were scaled from 0 to 360, and thresholding was applied based on the hue component to distinguish different colors within the stained collagen. Pixels with hue values between 330–359 and 0–20 were classified as red, 20–38 as orange, 39–61 as yellow, and 62–128 as green. The segmentation was applied to Insertion point and Free running regions of each sample. The proportion of each color was calculated as the number of pixels within each hue range, normalized to the total collagen content. The percentages of red, orange, and yellow pixels were summed to represent collagen type I, while the percentage of green pixels was reported as collagen type III.

## Results

### Protocol selection phase

Figure 2 presents mean ihMTR values calculated over the chosen ROIs (Figure 2-a) and the associated contrasts between structures (Figure 2-b) for different acquisition parameters during the Protocol selection phase on sample 1. Across all acquisitions, the ihMTR in free-running PF is consistently higher than in the myocardium and the Insertion point. Acquisition protocols 13 to 20 (see Table 2 for parameter values) are excluded from analysis due to their lower baseline ihMTR values in the free-running region (see Figure 2-a). Among the remaining acquisitions, we selected three protocols representing different configurations of saturation pulse duration and number of pulses per burst. The selected protocols, referred to as Protocols 1, 2 and 3 (Pr 1, Pr 2, and Pr 3 respectively) exhibited high ihMTR values in the free-running fiber (9.25%, 10.83%, and 9.33%, respectively in Figure 2-a), along with strong contrast both between free-running and myocardium (2.00%, 2.17%, and 2.07%, absolute ihMTR scale, in orange in Figure 2-b) and between free-running and the insertion point (3.40%, 3.39%, and 2.99% in purple in Figure 2-b).

**Figure 2:**
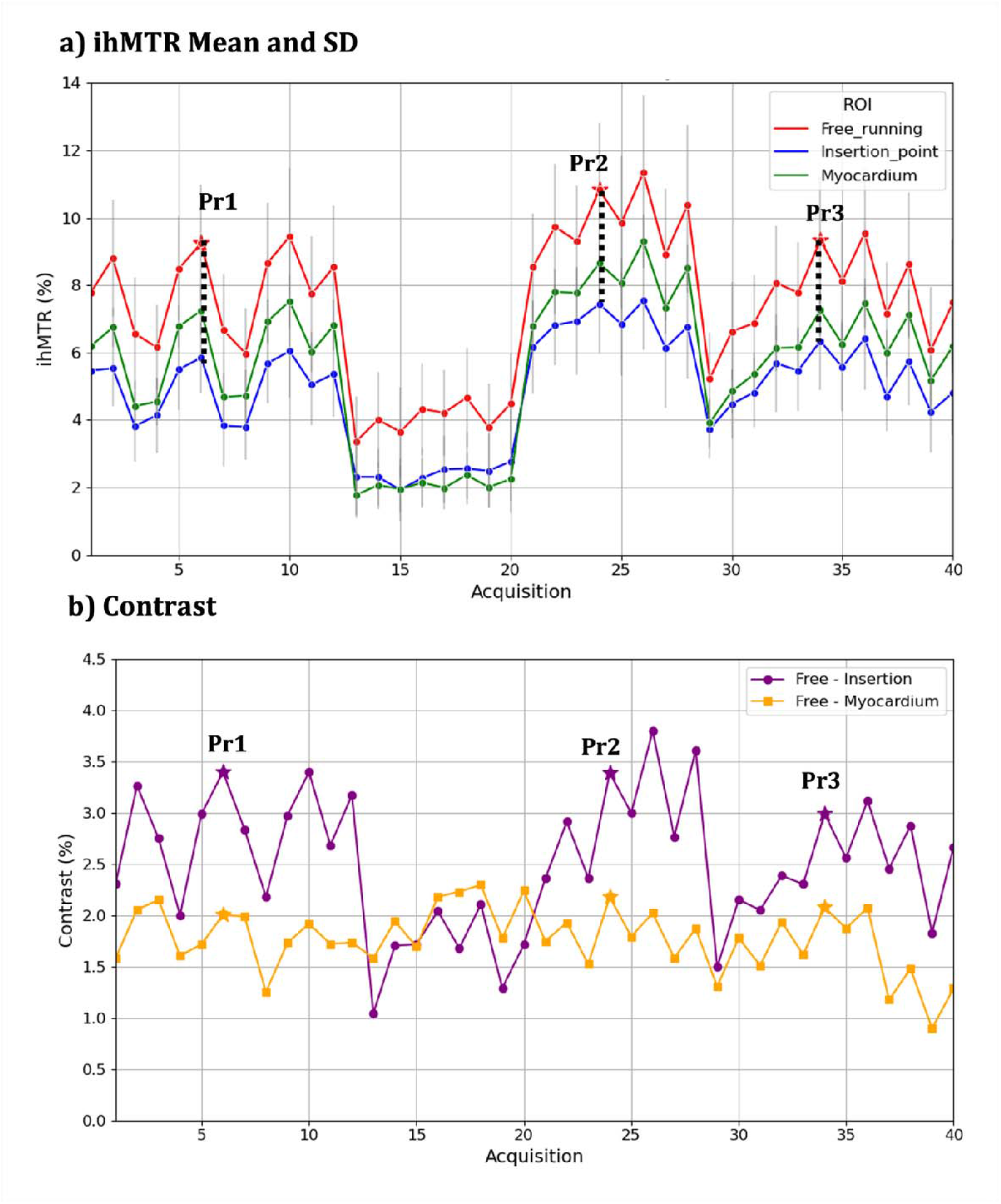
a) IhMTR in % for Free-running PF (red), Insertion point (blue) and myocardium (green) across all experiments. b) Contrast (in ihMTR units) for free-running PF vs. Myocardium (orange) and for free-running PF vs. Insertion point (purple) across all experiments. The three protocols selected for *Protocol application* phase are indicated on both figures.

### Protocol application phase

Figure 3 presents representative photographs of all samples used during the protocol application phase, together with the corresponding imaging slice, the ihMTR maps obtained using Protocol 1, and the selected ROIs in the free-running fibers (red), insertion points (blue), and myocardium (green). Overall, ihMTR values are higher in the free-running regions of the PF compared with the insertion points, whereas myocardial contrast exhibits substantial inter-sample variability.

**Figure 3:**
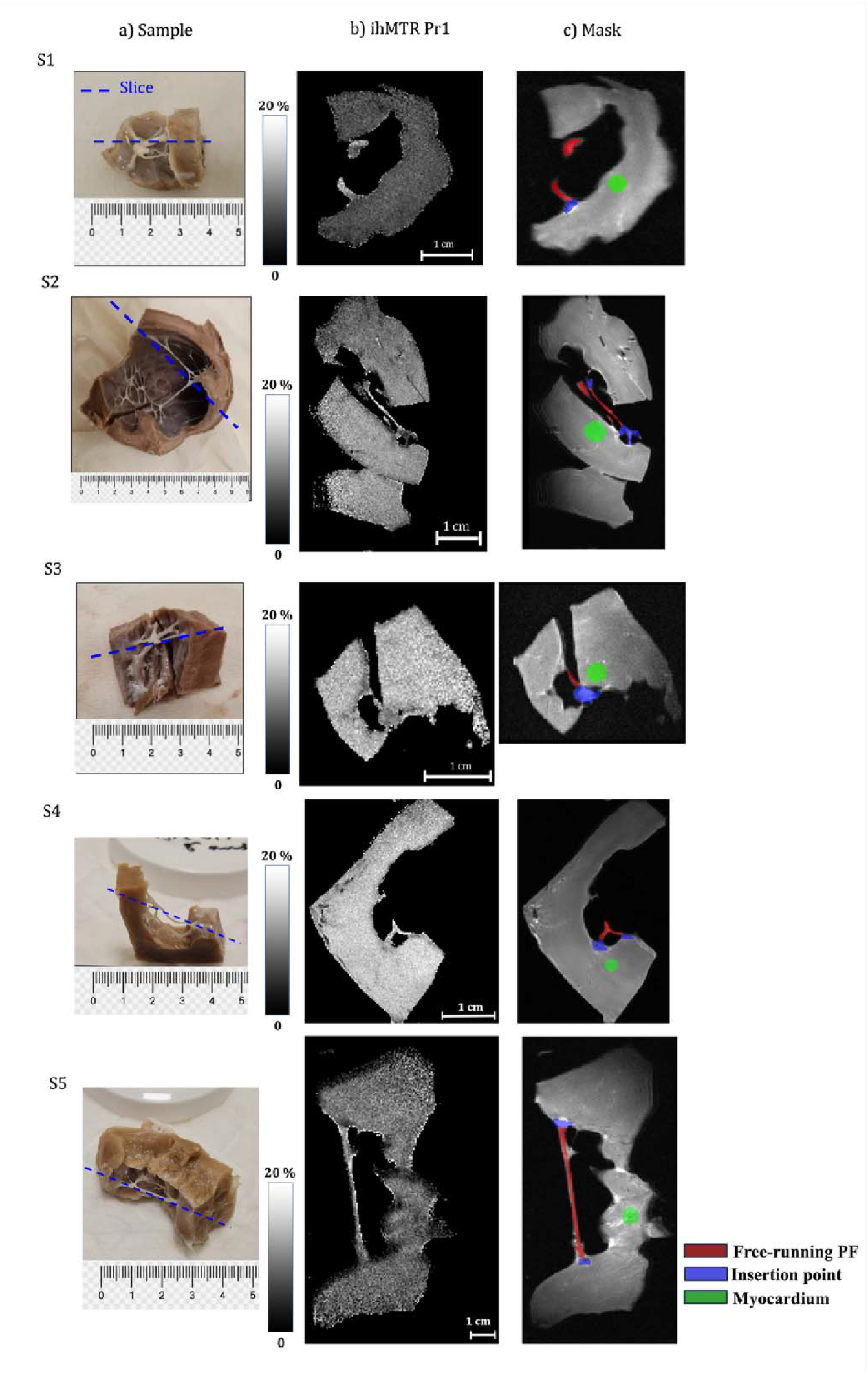
a) Photos of each sample with a dashed blue line indicating the image slice location (all in axial orientation) ; b) ihMTR from protocol 1; c) the mask used in the Protocol application phase differentiating the fiber’s insertion point (blue), the free-running section (red), and the myocardium (green) for each sample.

**Figure 4:**
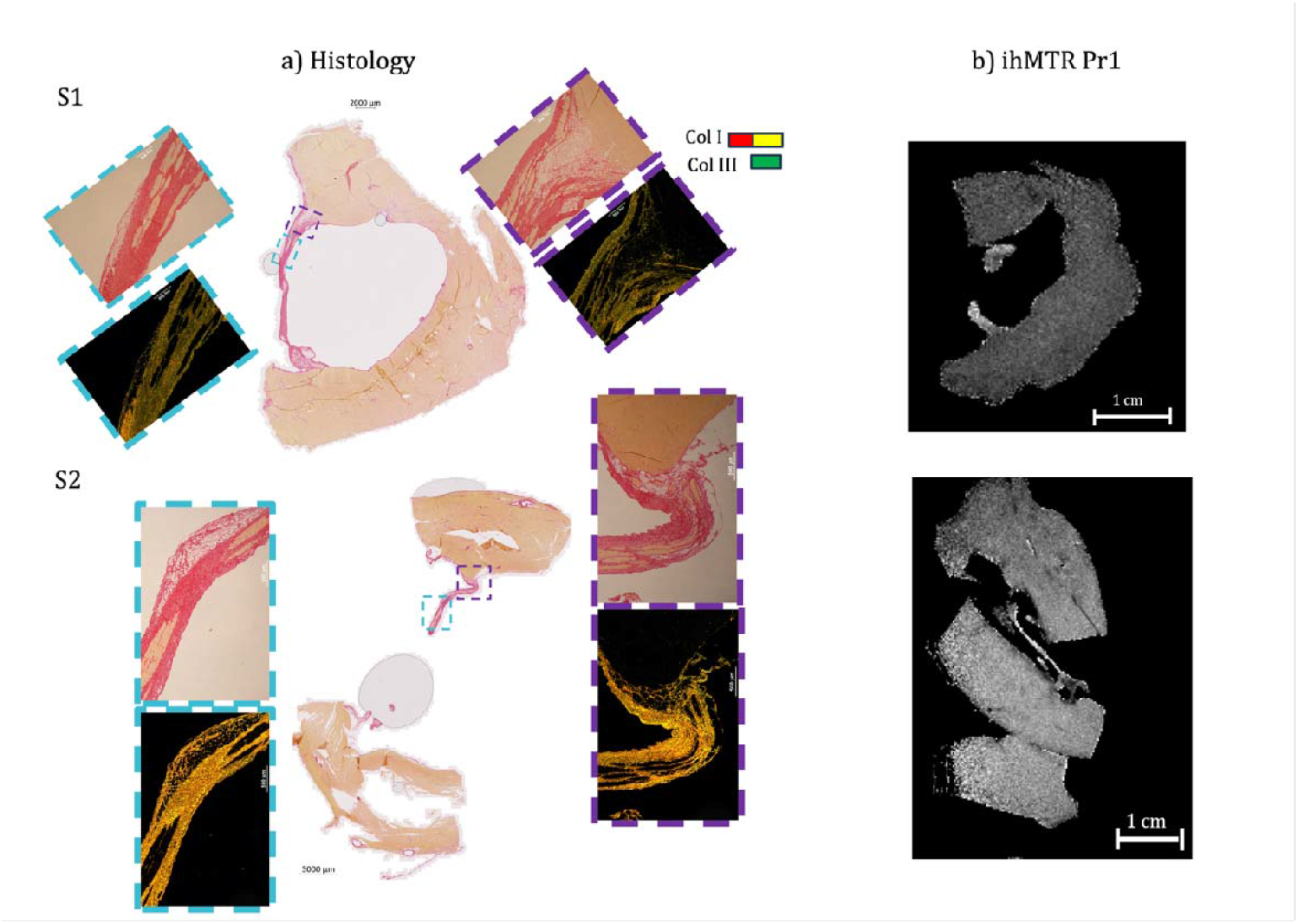
a) Histology images best matching the ihMTR image slice of two representative samples. Histological sections reveal structural details, including collagen, Purkinje cells, adipocytes, and myocytes. The polarized light (PL) filtered images depict collagen type I in red and yellow, while collagen type III appears in green. The light blue rectangle indicates the free-running and its corresponding zoomed-in images; the purple rectangle highlights additional zoomed-in views of the insertion point. b) The ihMTR map from Pr1 is also shown for comparison.

Detailed ihMTR mean ± SD values for each ROI, sample, and protocol are provided in Table S1 in Supplementary Materials for interested readers. Table 3 summarizes the ihMTR (mean ± inter-protocol SD), averaged across the three imaging protocols, for each ROI and for each of the five samples. Across all samples, ihMTR values are consistently higher in the free-running fiber region than at the insertion points (11.5 ± 1.5% vs. 9.0 ± 2.9%, respectively). In contrast, this trend was not observed in the myocardium: in samples S2 and S3, myocardial ihMTR values exceed those measured in the free-running regions (12.7 ± 1.2% vs. 11.9 ± 0.6% and 13.9 ± 1.1% vs. 11.0 ± 1.3% for myocardium vs. free-running regions in S2 and S3, respectively).

**Table 3:** ihMTR (Mean ± SD) averaged over 3 protocols in % for each ROI and each sample. The mean values are consistently higher in Free-running than in Insertion point although not statistically significant (Paired t-test p-value=0.6).

| #Sample | Free-running | Insertion point | Myocardium |
| --- | --- | --- | --- |
| 1 | 10.6±0.9 | 6.3±0.6 | 8.1±1 |
| 2 | 11.9±0.6 | 7.7±0.7 | 12.7±1.2 |
| 3 | 11±1.3 | 9.5±0.2 | 13.9±1.1 |
| 4 | 14.1±1.4 | 13.8±1.4 | 11±1 |
| 5 | 10.2±0.9 | 7.9±0.8 | 6.5±0.5 |

### Histological Analysis

Figure 3 presents histological sections (a) best matching the corresponding acquired ihMTR image (b) for two representative samples. The histological sections reveal the detailed architecture of the fibers, including Purkinje cells, collagen, adipocytes, and cardiomyocytes, supporting the identification of these regions as Purkinje fibers. Purkinje cells are identified by their distinctive shape and size, particularly in the free-running fiber region. Standard light microscopy and PL imaging provided complementary information about tissue organization, with collagen type I appearing in red, orange, or yellow, and collagen type III in green under PL.

PFs show a complex microstructure, with Purkinje cells surrounded by collagen (stained red under standard light) and adipocytes (visible as white aggregates). In each histological image, purple rectangles indicate zoomed-in views of the insertion point, highlighting the predominance of adipocytes in these regions. In contrast, light blue rectangles mark the free-running fiber regions, which are primarily enriched in collagen. These distinct structural patterns further support the differentiation between the insertion point and free-running subregions of the Purkinje network.

Table 4 presents the mean ± SD % of total collagen, adipocytes, collagen type I, and collagen type III for each sample, averaged across multiple high-magnification images acquired from the Free-running and Insertion point regions and across multiple microscope slides. A paired t-test over all samples revealed a significant difference in total collagen content between the Free-running and Insertion point ROIs (72.4 ± 15.9% vs. 31.1 ± 13.1%, respectively; p = 0.001). In contrast, adipocyte content is lower in the Free-running region than in the Insertion point, although this difference did not reach statistical significance. PL imaging indicates an overall predominance of collagen type I over collagen type III within the fibers (see Table 4). Collagen type I content is slightly higher in the Free-running region compared with the Insertion point (93.2 ± 3.1% vs. 85.7 ± 5.1%), whereas collagen type III content is modestly higher in the Insertion point than in the Free-running region (8.8 ± 3.9% vs. 3.0 ± 2.8%).

**Table 4:** Mean ± SD in % of total collagen, total adipocytes, collagen type I, and collagen type III for each sample, averaged across multiple images and microscope slides. The table reports both per-sample values and averages across all samples. A paired t-test indicates a significant difference in total collagen percentage between the Free-running and Insertion point ROIs. (p-value=0.001).

|  | % total Col |  | % Adp |  | % Col type I |  | % Col type III |  |
| --- | --- | --- | --- | --- | --- | --- | --- | --- |
| Sample | Free-running | Insertion point | Free-running | Insertion point | Free-running | Insertion point | Free-running | Insertion point |
| 1 | 84.3±12.3 | 44.7±19.2 | 5.7±3.8 | 17±7.9 | 93.5±2.1 | 87.3±6.3 | 2.2±2.3 | 8.6±6.4 |
| 2 | 77.7±9.3 | 45.5±3.4 | 1.8±4.9 | 8.5±4.9 | 96.4±1.1 | 93.1±1.4 | 0.4±0.7 | 3±1.3 |
| 3 | 87.7±7.3 | 25.1±14.2 | 0.1±0.1 | 20±8.2 | 95.7±1.2 | 80±18.2 | 0.5±0.4 | 9±5.9 |
| 4 | 49.4±13.6 | 16.8±5.2 | 6.4±3.2 | 11.4±7.3 | 88.7±5.3 | 86.2±4.7 | 7.1±6 | 9.4±5.3 |
| 5 | 62.9±7.5 | 23.4±7.2 | 5.1±5.1 | 4.8±2.6 | 91.7±2.6 | 81.7±7.9 | 4.6±2.9 | 14.1±8.2 |
| Mean±SD | 72.4±15.9 | 31.1±13.1 | 3.8±2.7 | 12.3±6.1 | 93.2±3.1 | 85.7±5.1 | 3.0±2.8 | 8.8±3.9 |

## Discussion

This study provides a proof of concept for the application of the inhomogeneous Magnetization Transfer (ihMT) technique to experimentally image the cardiac conduction system, specifically the Purkinje fibers. Imaging the His-Purkinje system presents significant challenges due to its fine anatomical structure, which requires high spatial resolution imaging and has distinct functional properties. To date, the most effective method for visualizing this network has been micro-CT with contrast agents applied to fixed cardiac samples ^8,10^. While micro-CT offers high spatial resolution and excellent visualization, its main limitation lies in the extensive sample preparation required, typically involving fixation, dehydration, and staining with contrast agents, which can damage tissue and prevent subsequent histological analysis, and in addition, is not transferable to in vivo imaging.

Our study demonstrates that ihMT offers a unique contrast mechanism for imaging Purkinje fibers, revealing their microstructural composition. By testing various ihMT saturation configurations on one sample, we identified three protocols that yielded the highest signal in fibers as well as high contrast between different regions of fiber and myocardium. In general, short pulse repetition times (Δt), high B_1_ saturation power and frequency offset in the order of 18 and 20 kHz are more suitable for imaging cardiac tissues. Conversely, long Δt values (3 ms) produced low ihMTR signals. This is expected from the theory of ihMT, and is attributed to the filtering of short T_1D_ components, as well as the amplitude loss of signal coming from longer T_1D_s ^33^. Increasing B_1_ should in principle increase ihMTR values. However, even though Specific Absorption Rate (SAR) regulations are less of a constraint in preclinical imaging as compared to human MRI, the hardware itself (power amplifier, RF coils, etc.) may become a limiting factor when the RF power approaches their tolerance limits, resulting in constraints in the maximal applied B_1_ RF excitation field strength that can be safely applied. In our study, we limited the B_1RMS_ to approximately 9.0 μT (with a maximum B_1peak_ at 46.4 μT), balancing safety requirements with the need for adequate signal excitation.

Notably, using the three optimized protocols, we identified that ihMTR signal varied along the length of the Purkinje fibers depending on anatomical location, whether in free-running segments or at insertion points. These findings are consistent with previously reported structural distinctions between the two regions ^7,34^. The free-running fibers, often referred to as false tendons, are composed primarily of conductive Purkinje cells surrounded by collagen and are specialized for rapid electrical conduction. In contrast, the insertion points, or Purkinje-myocardial junctions, are the terminal sites where impulses are transmitted from the Purkinje fibers to the slower-conducting working myocardium ^7^. These regions may also include an intermediate layer of transitional cells that facilitate signal propagation between the two tissue types ^34^.

### Effect of fiber composition

This heterogeneity was further confirmed by histological analysis, which revealed distinct variations in tissue composition along the fibers. Specifically, the insertion points exhibited a higher proportion of adipocytes, whereas the free-running segments contained greater amounts of collagen. The elevated ihMTR signal in the free-running fibers may thus be attributed to the presence of highly ordered collagen fibers, whose organized triple-helix structure is thought to enhance MT effects. In contrast, the lower ihMTR observed at the insertion points is likely associated with the lower total collagen content in this region. Additionally, the predominance of adipocytes at the insertion points likely contributes to the reduced ihMTR signal, as adipose tissue produces negligible magnetization transfer effects ^35^. Moreover, slightly higher levels of collagen type III were observed in the insertion point region, which may contribute to the observed contrast.

It is also important to note that the molecular composition of Purkinje fibers varies significantly across species, including sheep, pigs, rabbits, and humans ^36,37^. For instance, the study by Magat et al. ^14^ on porcine samples reported a markedly lower presence of adipocytes and a higher proportion of collagen type III in the insertion points of Purkinje fibers compared to the sheep samples examined in the present study. Additionally, Purkinje fibers can differ across species in terms of cell size and shape, as well as the amount and organization of surrounding connective tissue ^37^. These species-specific differences in fiber composition should be carefully considered when extrapolating findings to human physiology and clinical applications.

### Effect of fixation and storage

Interestingly, the average ihMT ratio in the myocardium showed considerable variability among the samples analyzed, in contrast to the fiber region, which exhibited less variability and better reproducibility. This disparity could arise from multiple factors. For instance, tissue fixation, which is known to alter water diffusion properties of biological tissues ^38^, may have contributed to these differences. While the fixation protocol was standardized for all samples, variations in the time elapsed between euthanasia and fixation, as well as the duration of the fixation process, could have impacted tissue properties. Additionally, changes in tissue characteristics over prolonged storage periods post-fixation have been reported in the literature ^39^, potentially affecting the experimental results. Finally, inter-sample variability could reflect intrinsic biological differences among the sheep. It may also result from samples with free-running fibers being taken from slightly different regions of the left ventricle.

### Effect of temperature and orientation

Temperature variations have a significant effect on ihMTR values within the PF, with an estimated increase of approximately 0.29% in absolute ihMTR scale per 1°C rise in temperature (Fig. S2). In contrast, the myocardium exhibited no significant changes in ihMTR with temperature variation, suggesting a tissue-specific sensitivity to thermal conditions. In the experiments in Protocol application phase, the temperature was maintained ± 1°C around 36°C, hence limiting ihMTR variations to ± 0.3% (absolute value), thereby ruling out the observed ihMTR differences being due to a temperature effect.

Additionally, the orientation of the fibers relative to the main magnetic field (B□) did not significantly influence ihMTR values in either the PF or the myocardium (Table S1). This indicates that, under our experimental conditions, fiber orientation is not a major confounding factor in the quantification of ihMTR.

## Conclusion

Despite the limited sample size and the use of 2D single-slice imaging, this ex-vivo high-resolution study demonstrates the potential of ihMT as a novel contrast mechanism to investigate the microstructural composition of Purkinje fibers. The findings suggest that ihMT may reflect the subtle differences in tissue structure, including the distribution of adipocytes, collagen, purkinje cells and cardiomyocytes, along the fibers. Moreover, this study lays the groundwork for developing biophysical models to interpret ihMT signals in terms of tissue composition and microstructure, providing a foundation for future studies aimed at quantifying tissue properties.

Future studies should aim to validate these findings on a larger sample size, explore 3D imaging techniques, and investigate the feasibility for in vivo applications of ihMT in cardiac imaging.

## Abbreviations

ihMT: inhomogeneous Magnetization Transfer
MT: Magnetization Transfer
PF: Purkinje fiber
CCS: Cardiac Conduction System
PL: Polarized Light
NEX: Number of excitations
SAR: Specific Absorption Rate

## Acknowledgement

This study received financial support from the French Government by the National Research Agency (ANR; SYNATRA ANR-21-CE19-0014-01) and Région Nouvelle Aquitaine (convention N°AAPR2022-2021-16609210). We would like to thank the preclinical platform team at IHU LIRYC, in particular Virginie Loyer and Coralie Lecompte, for their valuable assistance.

## References

1. Haissaguerre M, Vigmond E, Stuyvers B, Hocini M, Bernus O. Ventricular arrhythmias and the His–Purkinje system. Nat Rev Cardiol. 2016;13(3):155–166. doi:10.1038/nrcardio.2015.193

2. Atkinson A, Inada S, Li J, et al. Anatomical and molecular mapping of the left and right ventricular His-Purkinje conduction networks. J Mol Cell Cardiol. 2011;51(5):689–701. doi:10.1016/j.yjmcc.2011.05.020

3. Çetingül HE, Plank G, Trayanova NA, Vidal R. Estimation of Local Orientations in Fibrous Structures With Applications to the Purkinje System. IEEE Trans Biomed Eng. 2011;58(6):1762–1772. doi:10.1109/TBME.2011.2116119

4. Bogun F, Good E, Reich S, et al. Role of Purkinje Fibers in Post-Infarction Ventricular Tachycardia. J Am Coll Cardiol. 2006;48(12):2500–2507. doi:10.1016/j.jacc.2006.07.062

5. Garcia-Bustos V, Sebastian R, Izquierdo M, Molina P, Chorro FJ, Ruiz-Sauri A. A quantitative structural and morphometric analysis of the Purkinje network and the Purkinje–myocardial junctions in pig hearts. J Anat. 2017;230(5):664–678. doi:10.1111/joa.12594

6. Canale ED, Campbell GR, Smolich JJ, Campbell JH. Cardiac Muscle. Vol Vol. 2. Springer Science & Business Media; 2012.

7. Romero D, Camara O, Sachse F, Sebastian R. Analysis of microstructure of the cardiac conduction system based on three-dimensional confocal microscopy. PLoS One. 2016;11(10). doi:10.1371/journal.pone.0164093

8. Stephenson RS, Atkinson A, Kottas P, et al. High resolution 3-Dimensional imaging of the human cardiac conduction system from microanatomy to mathematical modeling. Sci Rep. 2017;7(1):7188. doi:10.1038/s41598-017-07694-8

9. Aminu AJ, Chen W, Yin Z, et al. Novel micro-computed tomography contrast agents to visualise the human cardiac conduction system and surrounding structures in hearts from normal, aged, and obese individuals: Iodine and graphene oxide – visualising human conduction system in normal, aged, and obese hearts. Translational Research in Anatomy.Elsevier GmbH. 2022;27. doi:10.1016/j.tria.2022.100175

10. Stephenson RS, Rowley-Nobel J, Jones CB, et al. Morphological Substrates for Atrial Arrhythmogenesis in a Heart With Atrioventricular Septal Defect. Front Physiol. 2018;9. doi:10.3389/fphys.2018.01071

11. Stephenson RS, Boyett MR, Hart G, et al. Contrast Enhanced Micro-Computed Tomography Resolves the 3-Dimensional Morphology of the Cardiac Conduction System in Mammalian Hearts. PLoS One. 2012;7(4):e35299. doi:10.1371/journal.pone.0035299

12. Chen W, Kuniewicz M, Aminu AJ, et al. High-resolution 3D visualization of human hearts with emphases on the cardiac conduction system components—a new platform for medical education, mix/virtual reality, computational simulation. Front Med (Lausanne). 2025;12. doi:10.3389/fmed.2025.1507005

13. Hwang M, Odening K, Ziv O, Choi BR, Koren G, Forder JR. Cardiac Purkinje Fiber Imaging: the first instance of in situ visualization of the conduction path using MR microscopy. Proc Intl Soc Mag Reson Med. Published online 2010.

14. Magat J, Fouillet A, Constantin M, et al. 3D magnetization transfer (MT) for the visualization of cardiac free-running Purkinje fibers: an ex vivo proof of concept. Magnetic Resonance Materials in Physics, Biology and Medicine. 2021;34(4):605–618. doi:10.1007/s10334-020-00905-w

15. Li Y, Casula V, Karjalainen J, Nissi MJ, Liimatainen T. Quantitative Magnetic Resonance Imaging of the Atrioventricular Conduction Axis Based on the Rotating Frame Relaxation Maps-Comparison with T2, T1 and Magnetization Transfer. Heart Rhythm. Published online June 2025. doi:10.1016/j.hrthm.2025.06.040

16. Laurent D, Wasvary J, Yin J, Rudin M, Pellas TC, O’byrne E. Quantitative and Qualitative Assessment of Articular Cartilage in the Goat Knee with Magnetization Transfer Imaging.; 2001. doi:10.1016/S0730-725X(01)00433-7

17. Wang C, Witschey W, Goldberg A, Elliott M, Borthakur A, Reddy R. Magnetization transfer ratio mapping of intervertebral disc degeneration. Magn Reson Med. 2010;64(5):1520–1528. doi:10.1002/mrm.22533

18. Jiang K, Ferguson CM, Ebrahimi B, et al. Noninvasive assessment of renal fibrosis with magnetization transfer MR imaging: Validation and evaluation in murine renal artery stenosis. Radiology. 2017;283(1):77–86. doi:10.1148/radiol.2016160566

19. Fuchs BC, Wang H, Yang Y, et al. Molecular MRI of collagen to diagnose and stage liver fibrosis. J Hepatol. 2013;59(5):992–998. doi:10.1016/j.jhep.2013.06.026

20. López K, Neji R, Mukherjee RK, et al. Contrast-free high-resolution 3D magnetization transfer imaging for simultaneous myocardial scar and cardiac vein visualization. Magnetic Resonance Materials in Physics, Biology and Medicine. 2020;33(5):627–640. doi:10.1007/s10334-020-00833-9

21. Varma G, Duhamel G, De Bazelaire C, Alsop DC. Magnetization transfer from inhomogeneously broadened lines: A potential marker for myelin. Magn Reson Med. 2015;73(2):614–622. doi:10.1002/mrm.25174

22. Girard OM, Prevost VH, Varma G, Cozzone PJ, Alsop DC, Duhamel G. Magnetization transfer from inhomogeneously broadened lines (ihMT): Experimental optimization of saturation parameters for human brain imaging at 1.5 Tesla. Magn Reson Med. 2015;73(6):2111–2121. doi:10.1002/mrm.25330

23. Varma G, Girard OM, Prevost VH, Grant AK, Duhamel G, Alsop DC. Interpretation of magnetization transfer from inhomogeneously broadened lines (ihMT) in tissues as a dipolar order effect within motion restricted molecules. Journal of Magnetic Resonance. 2015;260:67–76. doi:10.1016/j.jmr.2015.08.024

24. Alsop DC, Ercan E, Girard OM, et al. Inhomogeneous magnetization transfer imaging: Concepts and directions for further development. NMR Biomed. Published online August 29, 2022. doi:10.1002/nbm.4808

25. Mchinda S, Varma G, Prevost VH, et al. Whole brain inhomogeneous magnetization transfer (ihMT) imaging: Sensitivity enhancement within a steady-state gradient echo sequence. Magn Reson Med. 2018;79(5):2607–2619. doi:10.1002/mrm.26907

26. Forodighasemabadi A, Baucher G, Soustelle L, et al. Spinal cord and brain tissue impairments as long-term effects of rugby practice? An exploratory study based on T1 and ihMTsat measures. Neuroimage Clin. 2022;35:103124. doi:10.1016/j.nicl.2022.103124

27. Duhamel G, Prevost VH, Cayre M, et al. Validating the sensitivity of inhomogeneous magnetization transfer (ihMT) MRI to myelin with fluorescence microscopy. Neuroimage. 2019;199(May):289–303. doi:10.1016/j.neuroimage.2019.05.061

28. Rasoanandrianina H, Demortière S, Trabelsi A, et al. Sensitivity of the Inhomogeneous Magnetization Transfer Imaging Technique to Spinal Cord Damage in Multiple Sclerosis. AJNR Am J Neuroradiol. 2020;41(5):929–937. doi:10.3174/ajnr.A6554

29. Cabanis P, Magat J, Rodriguez-Padilla J, et al. Cardiac structure discontinuities revealed by ex-vivo microstructural characterization. A focus on the basal inferoseptal left ventricle region. Journal of Cardiovascular Magnetic Resonance. 2023;25(1). doi:10.1186/s12968-023-00989-y

30. Yushkevich PA, Piven J, Hazlett HC, et al. User-guided 3D active contour segmentation of anatomical structures: Significantly improved efficiency and reliability. Neuroimage. 2006;31(3):1116–1128. doi:10.1016/j.neuroimage.2006.01.015

31. Junqueira LCU, Bignolas G, Brentani RR. Picrosirius Staining plus Polarization Microscopy, a Specific Method for Collagen Detection in Tissue Sections. Vol 11.; 1979.

32. Jorgensen AM, Varkey M, Gorkun A, et al. Bioprinted Skin Recapitulates Normal Collagen Remodeling in Full-Thickness Wounds. Tissue Eng Part A. 2020;26(9-10):512–526. doi:10.1089/ten.tea.2019.0319

33. Hertanu A, Soustelle L, Le Troter A, et al. T1D-weighted ihMT imaging – Part I. Isolation of long- and short-T1D components by T1D-filtering. Magn Reson Med. 2022;87(5):2313–2328. doi:10.1002/mrm.29139

34. Tranum-Jensen J, Wilde AA, Vermeulen JT, Janse MJ. Morphology of electrophysiologically identified junctions between Purkinje fibers and ventricular muscle in rabbit and pig hearts. Circ Res. 1991;69(2):429–437. doi:10.1161/01.RES.69.2.429

35. Ercan E, Varma G, Dimitrov IE, et al. Combining inhomogeneous magnetization transfer and multipoint Dixon acquisition: Potential utility and evaluation. Magn Reson Med. 2021;85(4):2136–2144. doi:10.1002/mrm.28571

36. Elbrønd VS, Thomsen MB, Isaksen JL, et al. Intramural Purkinje fibers facilitate rapid ventricular activation in the equine heart. Acta Physiologica. 2023;237(3). doi:10.1111/apha.13925

37. Parto P, Tadjalli M, Ghazi SR, Salamat MA. Distribution and structure of purkinje fibers in the heart of ostrich (struthio camelus) with the special references on the ultrastructure. Int J Zool. 2013;2013. doi:10.1155/2013/293643

38. Thelwall PE, Shepherd TM, Stanisz GJ, Blackband SJ. Effects of temperature and aldehyde fixation on tissue water diffusion properties, studied in an erythrocyte ghost tissue model. Magn Reson Med. 2006;56(2):282–289. doi:10.1002/mrm.20962

39. Lohr D, Terekhov M, Veit F, Schreiber LM. Longitudinal assessment of tissue properties and cardiac diffusion metrics of the ex vivo porcine heart at 7 T: Impact of continuous tissue fixation using formalin. NMR Biomed. 2020;33(7). doi:10.1002/nbm.4298

