## Supplementary material for "Structural Characterization of Cardiac Purkinje Fibers Using Inhomogeneous Magnetization Transfer (ihMT): A proof of Concept MRI-Histology Approach": Supp materials

**Supplementary Materials**

| #Sample | Pr | ihMTR Free-running | ihMTR Insertion point | ihMTR Myocardium |
| --- | --- | --- | --- | --- |
| 1 | 1 | 9.9±3.3 | 5.6±1.5 | 7.2±1.0 |
|  | 2 | 11.7±3.2 | 7.0±1.8 | 9.2±1.1 |
|  | 3 | 10.1±2.7 | 6.2±1.6 | 7.8±1.3 |
| 2 | 1 | 12.1±6.7 | 7.2±3.5 | 12.2±1.4 |
|  | 2 | 12.3±5.4 | 8.5±3.4 | 14.2±1.4 |
|  | 3 | 11.1±4.7 | 7.2±3.1 | 11.8±1.4 |
| 3 | 1 | 10.7±3.3 | 9.8±2.7 | 14.7±1.7 |
|  | 2 | 12.5±3.5 | 9.5±2.4 | 12.6±1.9 |
|  | 3 | 9.9±3.3 | 9.3±2.7 | 14.3±1.9 |
| 4 | 1 | 15.6±3.0 | 15.5±1.1 | 12.3±2.1 |
|  | 2 | 12.6±2.4 | 12.7±1.0 | 10.3±1.9 |
|  | 3 | 13.9±2.3 | 13.3±1.2 | 10.5±2.1 |
| 5 | 1 | 10.5±3.4 | 7.1±1.8 | 6.7±3.7 |
|  | 2 | 10.8±3.0 | 8.8±2.1 | 6.8±2.8 |
|  | 3 | 9.1±3.0 | 7.8±1.9 | 5.9±3.0 |

Table S 1:  ihMTR values (mean ± intra-ROI SD) in % for each ROI, protocol, and sample.

**Orientation effect on ihMT**

To investigate the effects of orientation on the ihMTR signal, S1 was positioned in two different orientations, parallel to B_0_ and perpendicular to B_0_. The ihMTR values were then compared between the two orientations focusing on differences observed in the free-running PF and Myocardium regions, separately.

Table S2 presents the mean ± SD of ihMTR values in free-running PF and myocardium for each of the three protocols across different orientations. A Wilcoxon signed-rank test revealed no significant differences between orientations for both tissues, indicating that orientation does not have a significant effect on ihMTR values under these experimental conditions.

**Temperature effect on ihMT**

To investigate the effect of temperature on the ihMT signal, a series of experiments was conducted where the sample was heated to three target temperatures: 29°C, 34°C, and 38°C. At each temperature, acquisitions were performed, and the ihMTR was calculated for the free-running PF and myocardium. Figure S1 presents the mean ± SD of ihMTR values as a function of temperature in both regions. A linear regression and a two-sided t-test were applied to evaluate the dependence of ihMTR on temperature. The results show a statistically significant effect in the PF, with an average ihMTR increase of approximately 0.29% per 1°C rise in temperature. In contrast, temperature changes did not significantly impact ihMTR values in the myocardium.

To validate the accuracy of the temperature sensor used for monitoring the Fluorinert temperature during ihMT acquisitions and to assess the actual tissue temperatures, we employed a non-invasive MR thermometry method based on the proton resonance frequency (PRF) shift technique ^1^. The acquisition was performed using a fast low-angle shot (FLASH) sequence with the following parameters: TR = 10/30 ms, TE = 4.54 ms, matrix size = 160 × 120, field of view = 40 × 30 mm, in-plane resolution = 0.25 mm/pixel, slice thickness = 1 mm, and bandwidth = 50 kHz. Temperature images were processed using the PRF shift method ^1–3^ and displayed in real time using ThermoGuide software (Image Guided Therapy, Pessac, France). Additional offline analysis was performed using custom MATLAB software (MathWorks, R2018b), where ROIs were manually defined to extract pixel-wise temperature values within the PF and myocardium.

Using this setup, we monitored the temperatures in the free-running PF, myocardium, and Fluorinert (via the temperature sensor) before and after a 15-minute ihMT sequence. Figure S2 presents scatter plots of the recorded temperatures at both time points. The mean ± SD values were 38.7 ± 2.6°C in free-running PF, 37.5 ± 0.8°C in myocardium, and 36.8 ± 0.03°C at the sensor before ihMT; and 38.9 ± 2.7°C, 37.6 ± 0.6°C, and 37.0 ± 0.04°C, respectively, after the sequence. These findings confirm the reliability of the sensor and show that the temperature in the tissue remained stable throughout the experiment, thereby ruling out any modifications of ihMTRs resulting from temperature variations.

**Tables**

|  | PF | | | Myocardium | | |
| --- | --- | --- | --- | --- | --- | --- |
|  | Pr1 | Pr2 | Pr3 | Pr1 | Pr2 | Pr3 |
| 1^st^ Orientation: parallel to B_0_ | 9.7±3.1 | 11.8±3.5 | 10.1±3.2 | 6.5±1.4 | 8.8±1 | 7.2±1.2 |
| 2^nd^ Orientation: perpendicular to B_0_ | 9.5±3.4 | 11.4±3.3 | 9.8±2.8 | 7.2±1 | 9.3±1.2 | 7.8±1.2 |

Table S 2: ihMTR (mean ± SD) in % in PF and Myocardium for each protocol for sample 1 oriented parallel and perpendicular to B_0_. Wilcoxon signed-rank test shows no significant differences between the ihMTR from the two different orientations.

**Figures**

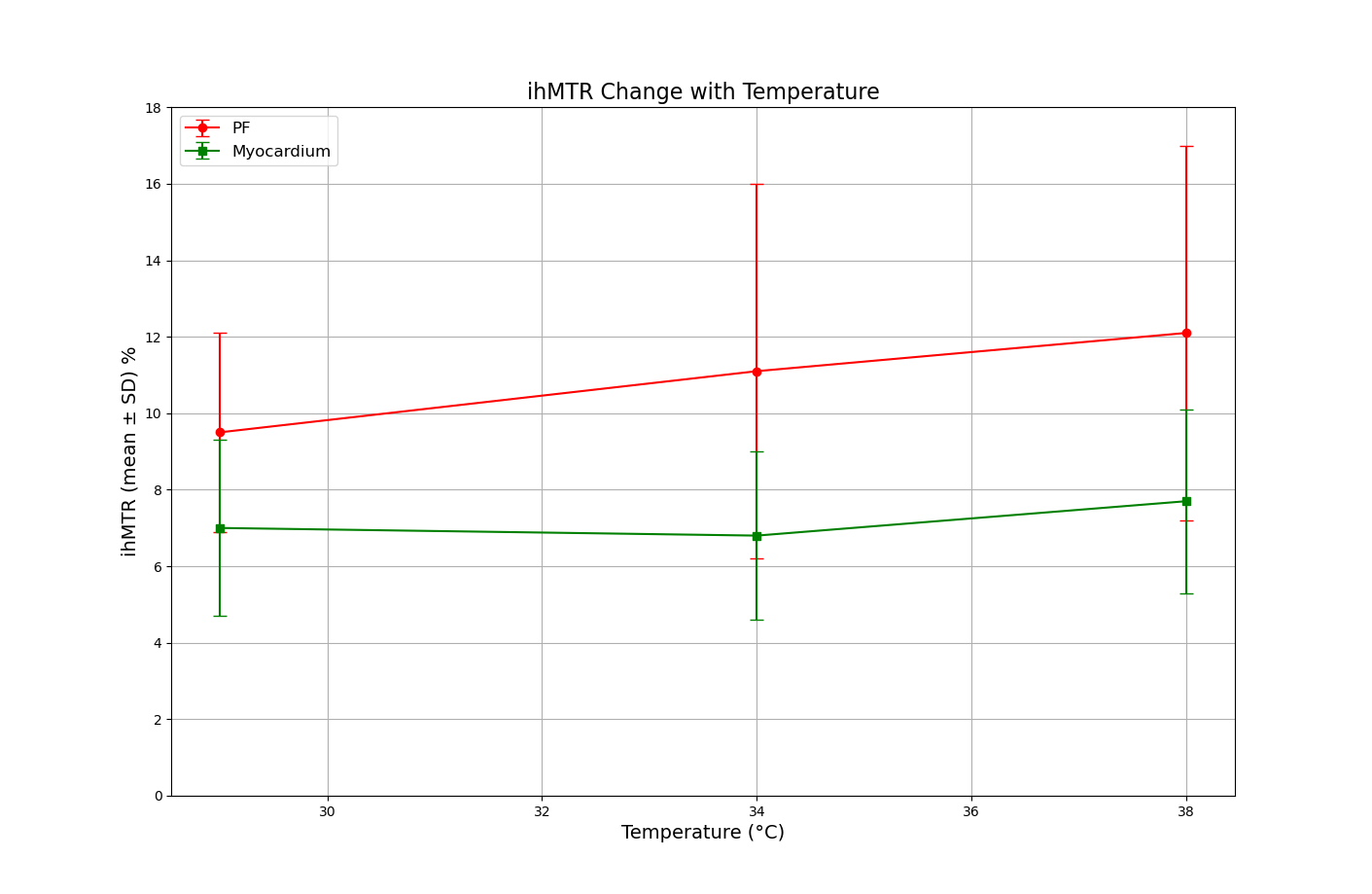

Figure S 1: Plot of mean ± SD of ihMTR in % as a function of temperature in PF (red) and Myocardium (green).

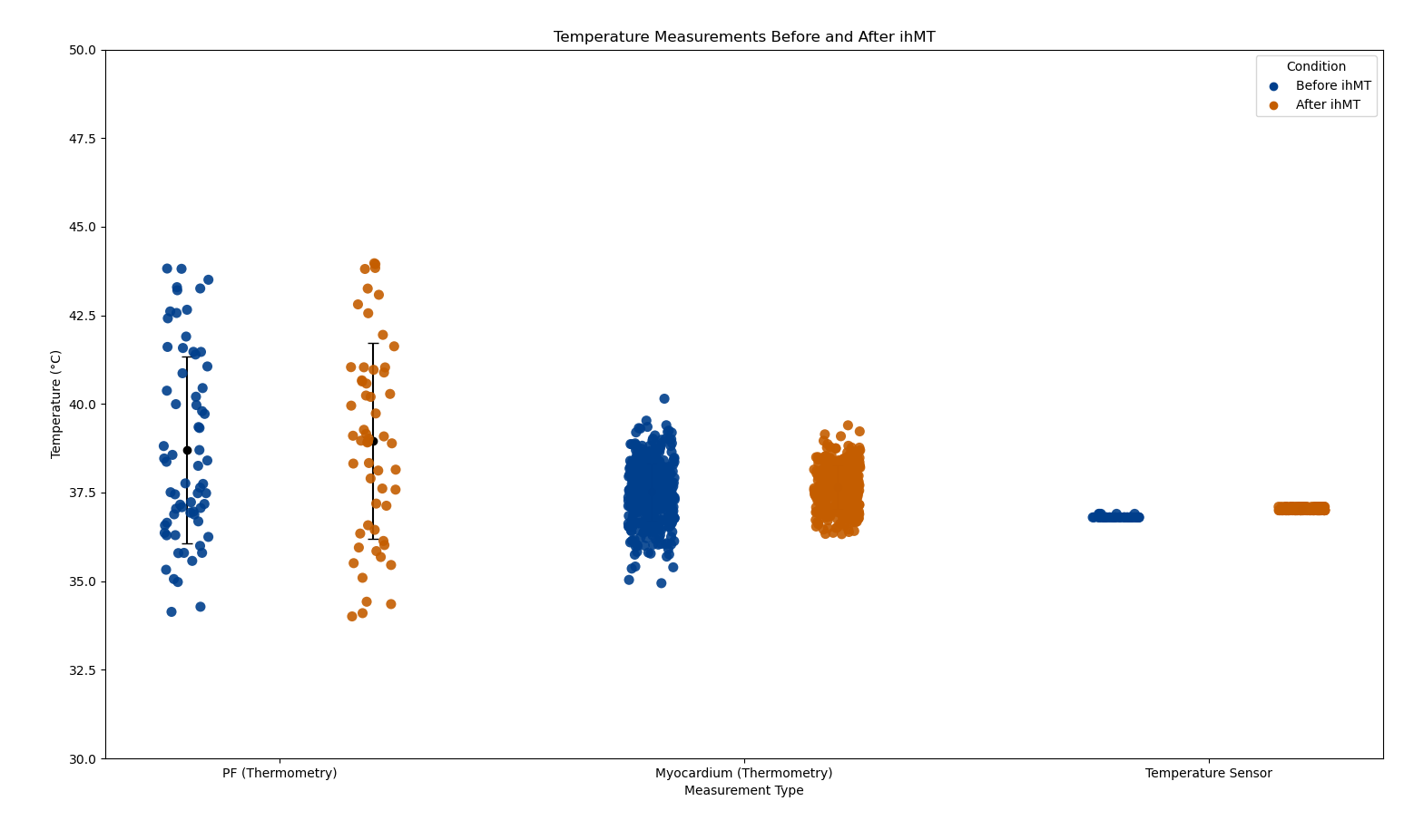

Figure S 2: Scatter plots showing the temperature in free-running PF and Myocardium evaluated from thermometry, as well as the temperature measured by the sensor, before and after the start of the ihMT sequence.

**References**

1.	De Poorter J, De Wagter C, De Deene Y, Thomsen C, Stahlberg F, Achten E. Noninvasive MRI Thermometry with the Proton Resonance Frequency (PRF) Method: In Vivo Results in Human Muscle. *Magn Reson Med*. Published online 1995. doi:10.1002/mrm.1910330111

2.	Ishihara Y, Calderon A, Watanabe H, et al. A Precise and Fast Temperature Mapping Using Water Proton Chemical Shift. *Magn Reson Med*. Published online 1995. doi:10.1002/mrm.1910340606

3.	Peters RD, Hinks RS, Henkelman RM. Ex Vivo Tissue-Type Independence Frequency Shift MR Thermometry. *Magn Reson Med*. Published online 1998. doi:10.1002/mrm.1910400316
